# Blind cavefish retain functional connectivity in the tectum despite loss of retinal input

**DOI:** 10.1101/2021.09.28.461408

**Authors:** Evan Lloyd, Brittnee McDole, Martin Privat, James B. Jaggard, Erik Duboué, German Sumbre, Alex Keene

## Abstract

Sensory systems display remarkable plasticity and are under strong evolutionary selection. The Mexican cavefish, *Astyanax mexicanus*, consists of eyed river-dwelling surface populations, and multiple independent cave populations which have converged on eye loss, providing the opportunity to examine the evolution of sensory circuits in response to environmental perturbation. Functional analysis across multiple transgenic populations expressing GCaMP6s showed that functional connectivity of the optic tectum largely did not differ between populations, except for the selective loss of negatively correlated activity within the cavefish tectum, suggesting positively correlated neural activity is resistant to an evolved loss of input from the retina. Further, analysis of surface-cave hybrid fish reveals that changes in the tectum are genetically distinct from those encoding eye-loss. Together, these findings uncover the independent evolution of multiple components of the visual system and establish the use of functional imaging in *A. mexicanus* to study neural circuit evolution.

## Introduction

Sensory systems are highly tuned through evolution to an animal’s environment and under stringent evolutionary pressure (*1, 2*). Adaptation to novel environments often results in modifications of sensory processing governing vital behaviors, including foraging and avoidance of predation (*3, 4*). The reliance on different sensory cues has been widely documented to be accompanied by changes in the anatomy of the primary sense organ, such as a tradeoff between expansion of the eye or olfactory epithelium, or changes in cortical area devoted to processing sensory information (*5–9*). However, little is known about the evolution of neural circuitry in downstream sensory processing centers.

The Mexican tetra, *Astyanax mexicanus* provides a powerful biological system to study the genetic and neural basis of sensory evolution, consisting of eyed surface populations that inhabit rivers in Northeast Mexico and southern Texas, and at least 30 cavefish populations (*7, 10*). Cavefish populations have evolved a series of morphological, physiological, and behavioral changes including eye loss, albinism, reduced sleep, loss of social behaviors, and expansion of non-visual sensory organs including olfaction, taste and mechanosensation (*11, 12*). Many of these phenotypes and behaviors have evolved repeatedly in geographically isolated cavefish populations, positioning *A. mexicanus* as a model to investigate the genetic and ecological factors that shape the evolution of traits (*13–15*). One of the most striking traits is convergence on eye loss, presumably to conserve energy in a perpetually dark environment (*16–18*). This phenotype has been studied for over 80 years, yet the concomitant physiological changes in visual processing centers have not been investigated.

In teleosts, the majority of retinal inputs project to the optic tectum, a brain region that is homologous to the mammalian superior colliculus (*19*). While the tectum primarily receives sensory inputs from the retina, minor inputs from auditory and lateral line modalities have been reported (*20*). In sighted species, visual processing within the tectum is essential for generating appropriate visuomotor behaviors including prey capture and predator avoidance (*21–23*). In comparison to surface fish, the size of the tectum is reduced in cavefish from multiple populations, revealing co-evolution of reduced inputs of the retina and its target centers within the brain (*6*). Recently, we generated a quantitative neuroanatomical atlas of surface fish and multiple cavefish populations that revealed that despite eye loss, the optic tectum is only reduced by ∼20% across populations, raising the possibility that this brain region has been repurposed for functions other than vision (*6, 24*). Therefore, the maintenance of the tectum despite the apparent loss of visual input provides a system to examine how brain structures are impacted by evolved differences in sensory input.

The optic tectum of the zebrafish, *Danio rerio*, is an established model to study development and function of visual sensory systems (*25, 26*). Two-photon Ca^2+^ imaging has been used to identify cell types that respond to different properties of visual stimuli, and infer the circuits functional connectivity (*27–30*). Recently, we have implemented transgenesis in *A. mexicanus*, permitting the application of approaches previously limited to zebrafish (*31*). The development of these tools has provided the first opportunity to investigate how neural responsiveness and functional connectivity of sensory circuits are shaped by evolutionarily derived changes in sensory function. Here, we used two-photon scanning microscopy to image surface and cavefish pan-neuronally GCaMP to directly define differences in neuronal dynamics within the tectum across independently evolution populations of *A. mexicanus*.

## Results

### Development of transgenic cavefish for brain imaging

Multiple populations of cavefish have converged on reduced eye size and volume of the optic tectum (*6, 16, 24, 32*), yet little is known about the underlying impacts on neuronal function. To investigate the changes in function of the optic tectum associated with eye loss we sought to compare the functional connectivity and the responsiveness to light in the tectum of river dwelling surface fish and two independently evolved populations of cavefish (Fig 1A). The Pachón population originates from old-surface stock and is located in the Sierra del Abra region, while the Molino population evolved from new surface stock and is located in the Sierra de Guatemala range (*33–35*). Geological and genomic evidence suggest these two populations of cavefish have independently evolved loss of eyes (*34–36*). To study changes in the function of the optic tectum in *A. mexicanus*, we generated surface and cave populations harboring the genetically encoded Ca^2+^ indicator (GCaMP6s) of the pan-neuronal promoter elavl3/HuC (*37*). The surface and Molino populations have previously been described (*6*); here, we also utilizea HuC:GCaMP6s Pachón line, allowing for direct comparison of brain function in independently evolved cavefish populations. Fish from the surface, Pachón, and Molino populations expressed the reporter as early as 1.5 dpf through 8.5 dpf (Fig 1B and Fig S1), confirming stable, brain-wide expression throughout early development across all three *A. mexicanus* populations. Therefore, these lines recapitulate expression in zebrafish and can be applied to measure neuronal activity throughout the brain.

**Figure 1:**
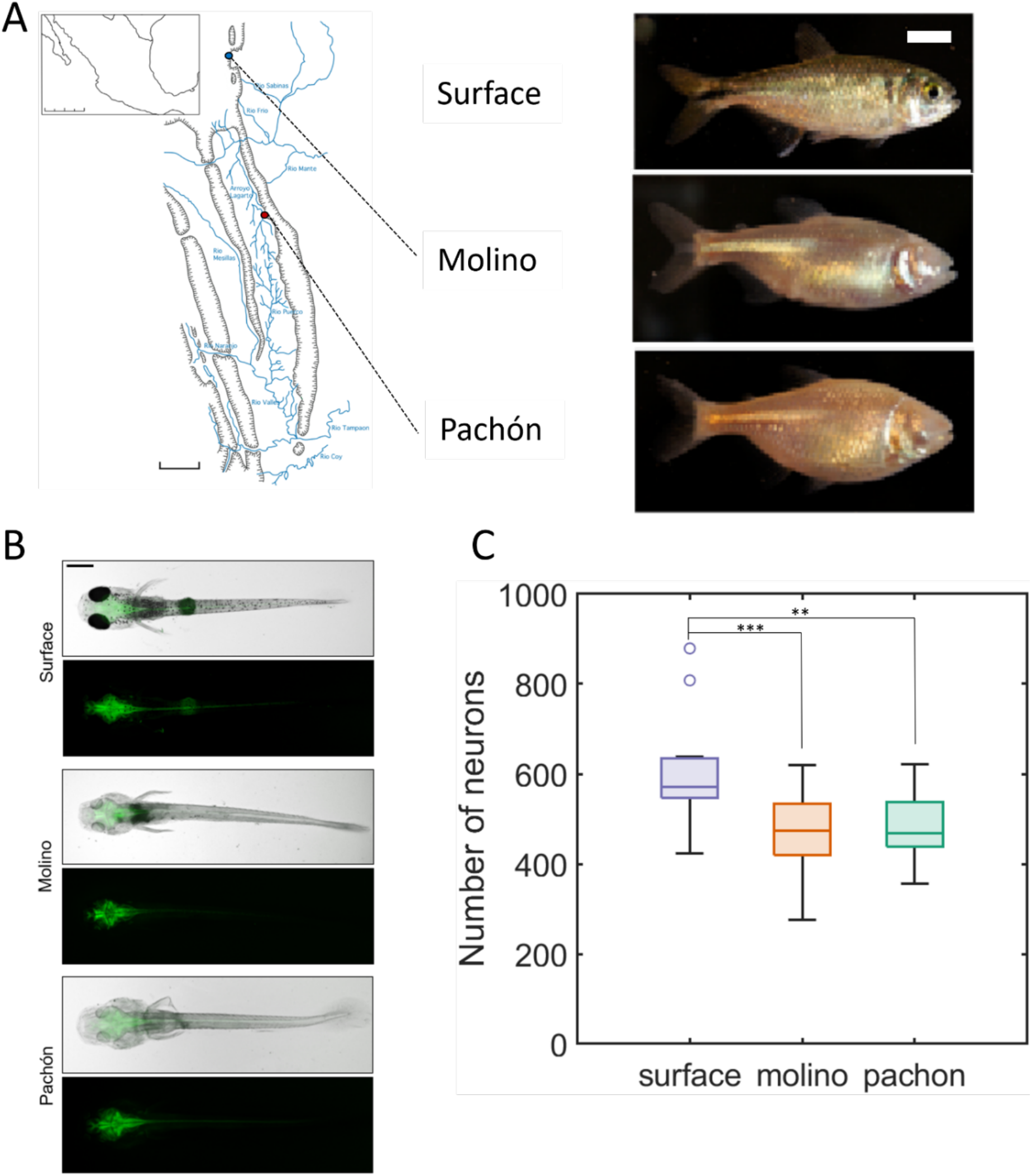
Generation of surface and cave fish transgenic lines expressing pan-neuronal HuC:GCaMP6s. **(A)** Left, a map of the Sierra del Abra region, identifying the location of the Pachón and Molino caves. Right, representative images of adult fish from the surface environment (top), and from the Molino (middle) and Pachón (bottom) caves. Scale bar = 0.5 cm. **(B)** Representative images of 6dpf surface, Molino, and Pachón larvae, demonstrating pan-neuronal expression (Top: bright-field image; bottom: fluorescent image). Scale bar = 0.5 mm. **(C)** Quantification of number of neurons in the tectum at the level of imaging, showing a reduction in tectal neuronal number in Molino and Pachón cavefish with respect to surface fish.

### Ongoing spontaneous activity in the optic tectum

Correlation in the spontaneous activity between neurons has been used as a proxy of the circuit’s functional connectivity. In zebrafish, the ongoing spontaneous activity of the optic tectum is organized in assemblies composed of highly correlated neurons with similar spatial tuning curves. These neuronal assemblies show attractor-like dynamics and may improve the detection of prey-like visual stimuli (*29, 30*). To quantify changes in the tectal functional circuitry that co-evolve with eye degeneration, we compared spontaneous activity in the tectum between surface and cave populations. For this purpose, we performed two-photon Ca^2+^ imaging of surface and cave transgenic GCaMP6s larvae (6 dpf) for a period of 30 minutes in the absence of any sensory stimuli (Fig S2). Quantification of the number of neurons in the tectum showed a reduction of ∼20% in the cavefish population, consistent with previous reports of reduced cavefish tectum size (Fig 1C), (*6*). We then analyzed the resulting fluorescence traces to detect significant Ca^2+^ transients as previously described (*38*), (Fig 2A-C). We first quantified the average frequency and duration of individual Ca^2+^ events across the tectal circuit. No differences were detected between surface fish and Pachón or Molino cavefish (Fig S3). Therefore, the overall levels of spontaneous activity within the tectum do not show evolutionary changes in two independently evolved cave populations. These findings reveal that the overall levels of spontaneous neuronal activity in the tectum are unaffected by evolutionary eye loss.

**Figure 2:**
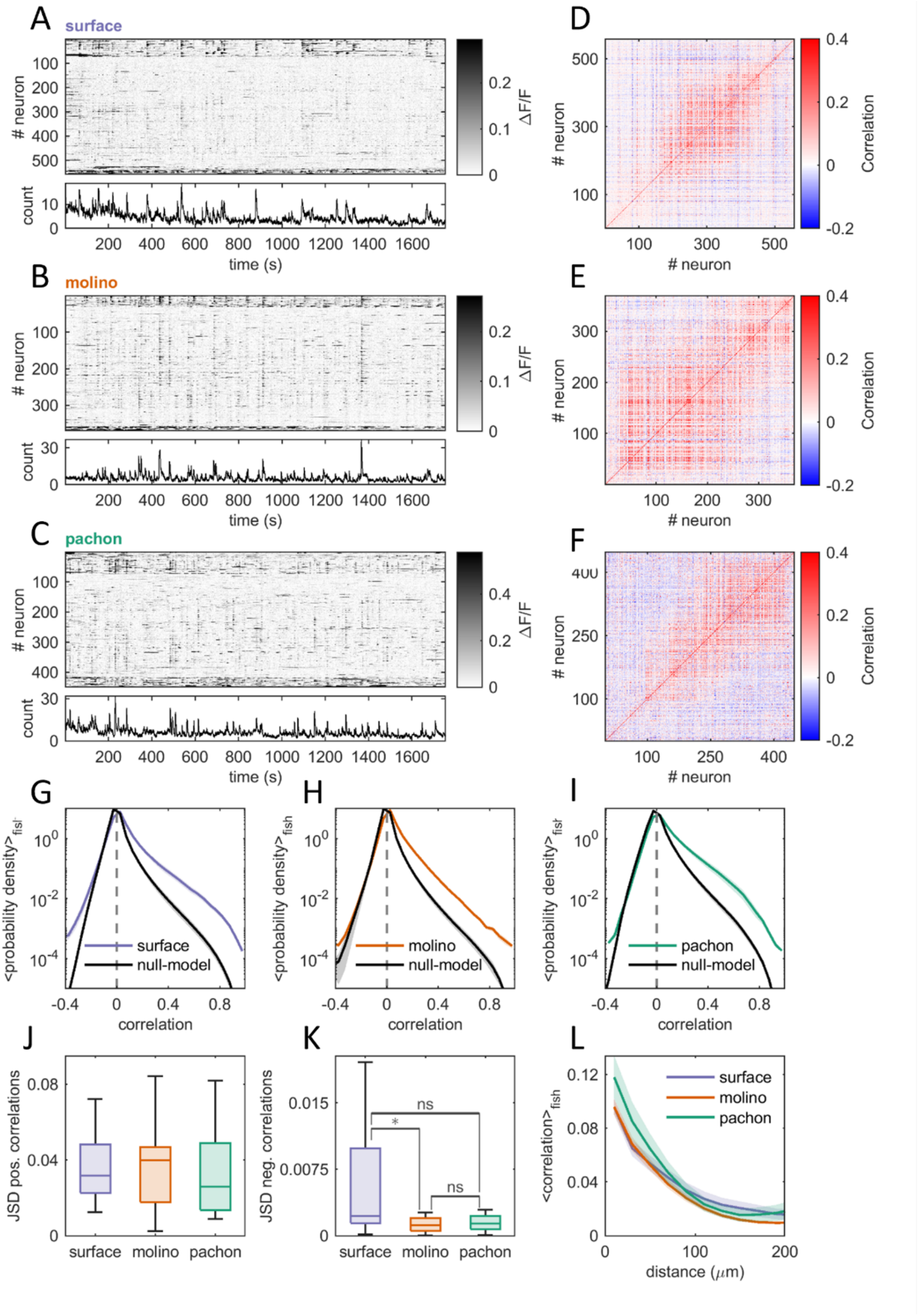
The structure of spontaneous activity is conserved between surface and cavefish. **(A-C)** Top: examples of raster plots of spontaneous neuronal activity recorded during 30 minutes in single surface (A), Molino (B), and Pachón (C) larvae, at a frame rate of 3.96 Hz. Color bar: relative change of fluorescence ΔF/F. The raster plots were sorted using the k-means algorithms with 10 clusters and Euclidean distance as a measure of similarity between neurons. Bottom: proportion of significant calcium events. **(D-F)** Example correlation matrices in single larvae corresponding to the raster plots in (A,C,E). Blue: negative correlations, red: positive correlations. The correlation matrices were not sorted according to the k-means clustering solution used in (A,C,E). **(G**,**H**,**I)** Probability density distribution of pair-wise spontaneous neuronal activity correlations across larvae in surface (G, blue curve, n=12), Molino (H, orange curve, n=20) and Pachón (I, green curve, n=17). In black is the average of the same distributions after circularly shifting each neuron’s calcium activity time series by a random lag (between 0 and 30 minutes). The ensures that the temporal structure of each time series is preserved while pair-wise correlations are destroyed (null-model). Shaded area: mean ± standard deviation. **(J,K)** For each larva we measured the probability density function of pair-wise spontaneous correlations and quantified the difference with the null-model by computing the Jensen-Shannon divergence (JSD) between the two distributions for positive (J) (Surface: 0.035 ± 0.018 JSD, Molino: 0.036 ± 0.021 JSD, Pachón: 0.037 ± 0.030 JSD, p=0.84; Kruskal-Wallis non-parametric ANOVA) and negative (K) (Surface: 0.0062 ± 0.0073 JSD, Molino: 0.001 ± 0.0020 JSD, Pachón: 0.002 ± 0.0047 JSD, p=0.052; Kruskal-Wallis non-parametric ANOVA) correlations. Statistical significance was assessed at the group level using the non-parametric Kruskal-Wallis one-way analysis of variance (p=0.841 for positive correlations, p=0.0525 for negative correlations), followed by post-hoc testing using the Tukey-Kramer procedure between each pair (* : p<0.05, ns : not significant). Blue: surface, n=12; orange: Molino, n=20; green: Pachón, n=17. **(L)** Average distribution of pair-wise correlations against distance across larvae. For each pair of neurons, we measured the pair-wise correlation coefficient as well as the physical distance in the optical section between those two neurons in micrometers. We averaged the pairwise correlations within distance bins across larvae in surface (blue, n=12), Molino (orange, n=20), and Pachón (green, n=17). Shaded gray area: mean ± standard deviation.

To study the effects of evolutionary eye loss on the functional connectivity of the optic tectum, we analyzed the tectal spontaneous activity in surface and cavefish.

To determine whether correlated activity represents statistically significant interactions that are indicative of neural circuit connectivity in the optic tectum, we applied Pearson’s pairwise correlations between the spontaneous activity of the tectal neurons. The probability distribution of the correlations in spontaneous activity were compared with a null model that randomly shifted the time series of every neuron. To quantify the difference between the two distributions of the pair-wise correlations, we used the Jensen-Shannon Divergence (JSD), which measures the similarity between two probability distributions (see methods), (Fig 2G-I). We found no significant differences in the pair-wise positive correlations between the tectal neurons of surface fish, or Pachón and Molino cavefish (Fig 2J). These findings suggest that functional tectal connectivity was retained in cavefish despite eye loss. However, analysis of the negative correlations revealed a significant reduction in the Molino cavefish, with Pachón approaching significance (Fig 2K). Given the importance of the inhibitory/excitatory balance to visual processing, the reduction in negative correlations in cavefish may reflect a physiological correlate of relaxed selection on the visual circuit in blind cavefish that is driven by a lack of retinal input. Finally, analysis of the physical distance between correlated neurons revealed that both surface and cave populations exhibit spatially structured correlations, with an exponential decrease in correlations with the Euclidean distance between neurons (Fig 2L). Therefore, positive correlations are resistant to the evolved loss of visual function, suggesting that excitatory and inhibitory connectivity are under partially independent evolutionary selection.

### Visual responses in the optic tectum

In zebrafish and other teleosts, the optic tectum is the main visual processing center (*25, 39, 40*). Cavefish retain a diminished eye, and they behaviorally respond to some light cues; however, the eye is suggested to be non-functional as cavefish exhibit severely reduced retinotectal connections by adulthood (*41*). To examine whether cavefish retain a light-evoked response in the optic tectum, we used two-photon microscopy to image light-evoked activity in transgenic GCaMP6s surface and cavefish (6 dpf). Briefly, fish were immobilized in low-melting point agarose and imaged during visual stimulus presentation, which consisted of a white-light LED located in front of the larvae. The LED was turned on for 30 sec., and presented ∼ 10 times, separated by 60 sec. (see methods, Fig S4A, S5). Using a multivariate linear regression model we categorized the Ca^2+^ responses of individual tectal neurons during stimulus presentation and identified neurons with multiple distinct response profiles (Fig 3A). In surface fish, the most common were cells which responded to either the onset (“On” cells”) or offset (“Off” cells) of light, with another subset which responded to both the onset and offset of light (“On/Off” cells). In addition, we identified a small number of neurons that exhibited a sustained activation throughout the period of the light stimulus (“Sustained” cells), and another that showed a sustained decrease in activity during light stimulus (“Inhibited” cells). Therefore, the tectal light response in *A. mexicanus* surface fish contains cell types previously observed in zebrafish (*42, 43*).

**Figure 3:**
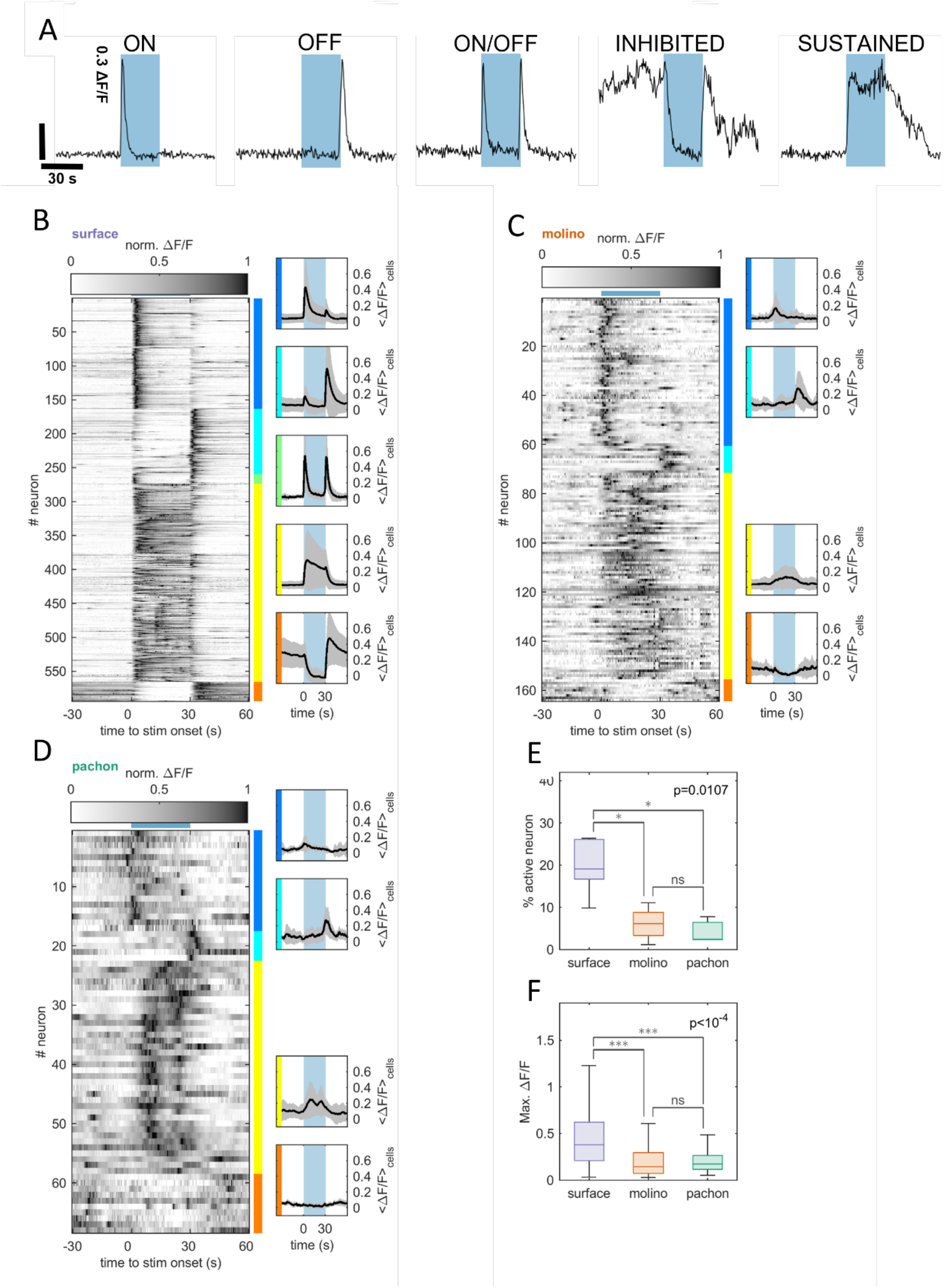
Partial loss of neuronal response to visual stimuli in cavefish populations. **(A)** Representative traces of the five types of light response. **(B-D)** Left: raster plots of normalized neuronal activity over the whole population of surface (A, n=6), Molino (B, n=5), and Pachón (C, n=3), averaged around the onset of whole-field visual stimulation. Visual stimuli were delivered by an array of white LEDs for a duration of 30 seconds (represented by horizontal light blue bar above the raster), with an inter-stimulus interval of 120 seconds, and neuronal activity was recorded with a frame rate of 1.96 Hz. Each experiment contains between 8 and 20 visual stimuli. Neurons responsive to the visual stimuli were detected using a linear regression model and classified into 5 categories: “on” cells (blue), “off” cells (turquoise), “on/off” cells (green), “sustained” cells (yellow) and “inhibited” cells (orange). Right panels: ΔF/F averaged across neurons and stimuli repetitions for each response type. Average slope of the response onset for surface: 0.106 ± 0.008 ΔF/F.s^-1^, Molino: 0.009 ± 0.005 ΔF/F.s^-1^, Pachón 0.007 ± 0.002 ΔF/F.s^-1^, R-squared surface: 0.826 ± 0.012, Molino: 0.853 ± 0.077, Pachón: 0.356 ± 0.197, p=0.008 Kruskal-Wallis). Shaded gray area: mean ± standard deviation. **(E)** Proportion of neurons responding to the whole-field visual stimulus in one hemi-tectum. Surface: 19.55% ± 6.26%, Molino: 6.10% ± 3.78% and Pachón: 4.20% ± 3.10%, mean ± standard deviation, p=0.0107 Kruskal-Wallis non-parametric ANOVA. Statistical significance was assessed at the group level using the non-parametric Kruskal-Wallis one-way analysis of variance (p=0.0107), followed by post-hoc testing using the Tukey-Kramer procedure between each pair (* : p<0.05, ns : not significant). **(F)** Distribution of the peak values for the amplitude of the neuronal response (ΔF/F) to whole-field visual stimuli. (Surface: 0.491 ± 0.432 ΔF/F, Molino: 0.212 ± 0.192 ΔF/F and Pachón: 0.241 ± 0.212 ΔF/F, mean ± standard deviation, p<10^−4^ Kruskal-Wallis non-parametric ANOVA), Kruskal-Wallis one-way analysis of variance (p<10^−4^ for positive correlations, p=0.0525 for negative correlations), Tukey-Kramer post-hoc test (*** : p<0.001, ns : not significant).

While cavefish have previously been reported to be blind with the exception of response to looming stimulus and lack physiological response to light in the tectum (*44, 45*), we found that both populations of cavefish showed significant, although severely reduced, Ca^2+^ responses to the presented visual stimuli (Fig 3B-D). In comparison to surface fish, Pachón and Molino larvae showed a significantly smaller number of responsive neurons to the visual stimulus, representing a reduction of responding neurons of approximately 75% in Molino, 90% in Pachón (Fig 3E). The amplitude of the Ca^2+^ responses of the cavefish to the visual stimulus was also reduced by ∼60% (Fig 3F). The probability of the neurons to respond to visual stimuli was also significantly smaller in cavefish with respect to surface fish (Fig S5). The onset precision of the visual response with respect to the onset of the stimulus was significantly more variable in the cavefish with respect to that of the surface fish. Moreover, the “On/Off” neurons were absent in both Pachón and Molino larvae, and the inhibited neurons showed very weak responses that were almost absent in Pachón (Fig 3B-D). Together, these findings reveal that although cavefish exhibit severe reductions in visual tectal function, they maintain the genetic and cellular functions necessary to process visual stimulus.

The presence of light responsiveness in the cavefish tectum raises the possibility that the retina is still functional in larval cavefish. Alternatively, responses in the tectum may be downstream of extra-retinal light-responsive organs such as the pineal gland (*46*). To distinguish between these possibilities, we imaged the retina response in GCaMP-expressing Molino and Pachón larvae while presenting a series of 30 sec. light stimuli. Due to pigmentation, recording of the intact retina in surface fish was not possible. We identified strong spontaneous retinal waves known to play a role in the development of retinotectal pathways (*47–49*). However, no light-evoked activity was detected in either cave population (Movie 1). These findings suggest that although the retina is still alive and active, it is non-functional, and light responsiveness in the cavefish tectum must stem from a non-visual source (e.g. the pineal gland or deep-brain internal photoreceptors) (*46, 50*).

### Analysis of Visual response in hybrid populations

It is possible that the decrease in the evoked tectal light response in cavefish is due to the partial loss of eyes that, at 6 dpf, are still in the evolutionary process of degenerating (*51*). The ability to generate surface-cave hybrid fish has been widely used to examine the functional and genetic relationship between traits (*52*). To determine whether changes in the anatomy of the tectum are related to the loss of eyes, we examined the relationship between traits in genetically and morphologically variable hybrid fish. By crossing transgenic HuC:GCaMP6s surface fish to Pachón or Molino cavefish and re-crossing the siblings for one generation, we generated genetically variable F2 surface x cave hybrids harboring the HuC:GCaMP6s transgene (Fig 4A). To determine whether there is a developmental or genetic relationship between eye size and the number of neurons in the tectum we examined the relationship between these two traits. We found an association between eye diameter and the number of neurons in the tectum in hybrids from the Molino, but not Pachón, caves (Figure 4B). These findings suggest that the reductions in the size of the tectum and eyes of Molino cavefish is controlled by shared genetic architecture, while these factors are independent in Pachón cavefish, suggesting that the evolutionary mechanisms that confer eye loss and/or tectal size in these two populations differ. The finding that tectal development is linked to eye size in Molino hybrids led us to question whether the tectum’s functional response is also related to eye size. We measured eye size in F2 fish, performed two-photon Ca^2+^ imaging in response to changes in illumination, and examined correlations between the two. We found no significant relationship between eye size and tectal response, in any of the cell types previously identified (Figure S6). Therefore, there is a functional relationship between eye and tectum size in Molino, but eye size does not impact light-induced tectum activity in either cavefish population.

**Figure 4.**
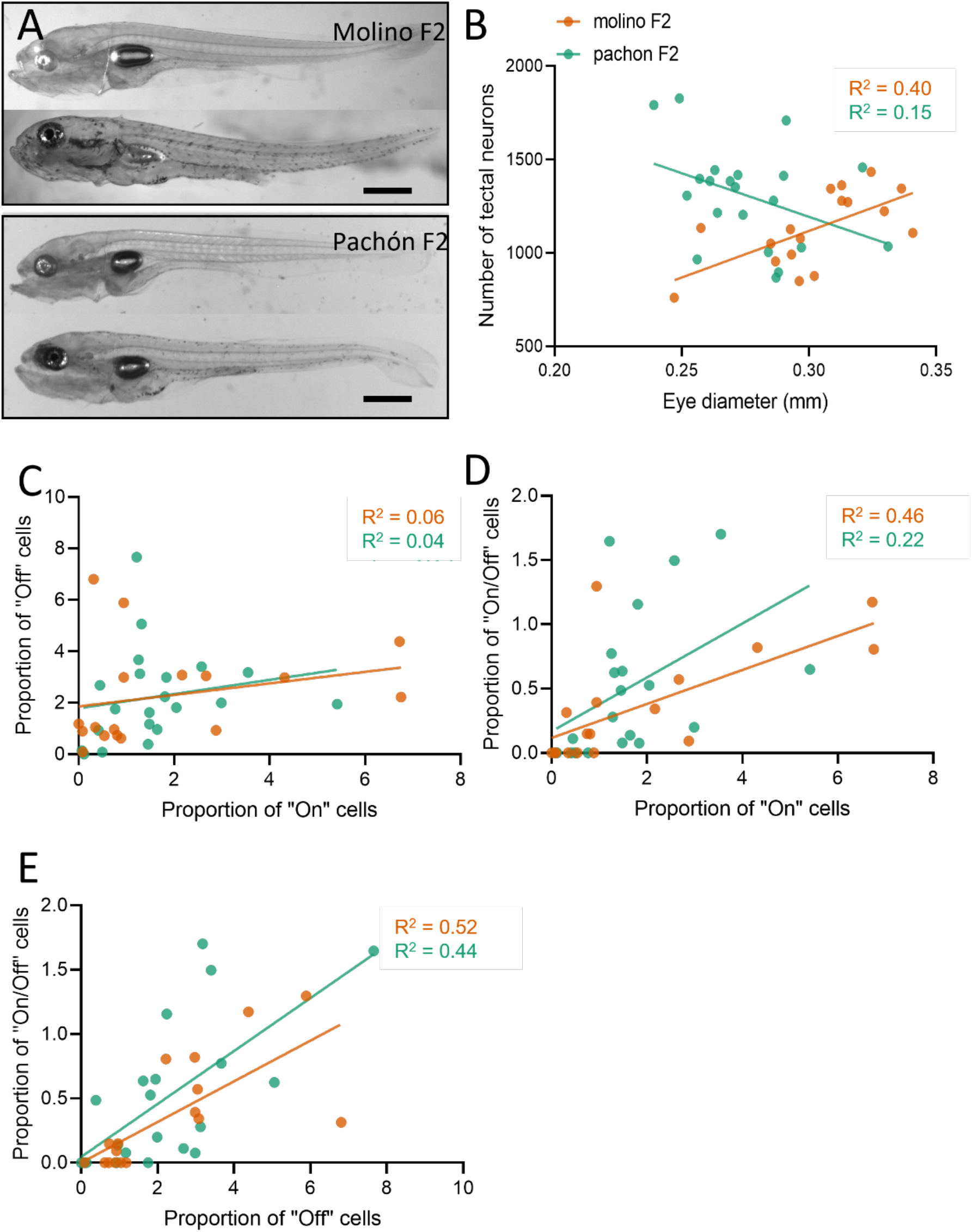
Analysis of light response in a hybrid F2 population. **(A)** Examples of 6dpf F2 crosses, demonstrating variability in eye size and pigmentation. **(B)** Molino F2s (orange) exhibit a correlation between eye size and tectal neuron number (*F*_1,15_ = 10.06, *p*=0.0063), while Pachón F2s (green) do not (*F*_1,19_ = 3.399, *p*=0.08). **(C)** There is no relationship between the proportion of “on” and “off” cells in the tectum of either Molino (*F*_1,15_ = 1.013, *p*=0.33) or Pachón (*F*_1,19_ = 0.711, *p*=0.41) F2s. **(D)** There is a significant correlation between the proportion of “on” and “on/off” cells in the tectum of both Molino (*F*_1,15_ = 12.56, *p*=0.003) and Pachón (*F*_1,19_ = 5.261, *p*=0.03) F2s. **(E)** There is a significant correlation between the proportion of “off” and “on/off” cells in the tectum of both Molino (*F*_1,15_= 16.03, *p*=0.001) and Pachón (*F*_1,19_= 14.97, *p*=0.001) F2s.

To determine whether there is a shared genetic basis underlying the development of distinct neuronal functional populations in the tectum, we quantified different neuronal response types in surface-cave hybrids. In both Pachón and Molino cavefish there was no relationship between the number of “On” cells and “Off” cells, suggesting that each population is independently regulated (Figure 4C). This finding suggests that distinct genetic pathways guide the development of these cell types within the tectum. However, a relationship was identified between each singly responsive cell type and dual-responsive “On/Off” cells, suggesting a shared genetic basis (Figure 4D, 4E). Additional correlation analyses between incidence of cell types, including the “Sustained” and “Inhibited” cells, yielded no significant correlations (data not shown). Taken together, these findings suggest shared genetic architecture underlies the loss of “On/Off” cells and cells that exclusively respond to light onset or offset across both Molino and Pachón populations of cavefish. These findings are indicative of convergence on shared developmental processes contributing to changes in tectum function across independently evolved cavefish populations, through different genetic mechanisms.

## Discussion

Cave animals have evolved numerous phenotypic changes that are likely adaptive to a cave environment, including eye loss and expansion of olfactory and mechanoreceptive organs (*12, 53, 54*). These changes occur in diverse cavefish species throughout the world, suggesting the shared ecological features in cave environments, such as perpetual darkness and reduced food availability, underlie the evolved changes in sensory processing (*55*). The repeated evolution of eye loss in the different *A. mexicanus* cavefish populations has provided a model to investigate the genetic basis of evolution. Several studies have identified numerous factors underlying eye loss including a critical role for the degeneration of the lens and retina (*56–60*). Further, genetic complementary studies have revealed that different genetic mechanisms underlie eye loss across cave populations (*36, 61*). Despite our detailed understanding of genetic and morphological changes that occur in sensory organs, strikingly little is known about evolutionary adaptations in sensory processing centers of the brain. We utilized GCaMP-expressing transgenic *A. mexicanus* to examine the changes in the main visual processing center in surface and cavefish, the optic tectum.

In teleost fish, the optic tectum is the main visual processing region, it is involved in the detection and processing of visual stimuli to generate goal-directed motor behaviors such as prey capture (*62*). Similar to our findings in *A. mexicanus*, numerous cellular response types have been previously identified in the zebrafish tectum, including those which respond selectively to changes in light level (*63*). We identified shared cellular response types across surface and both cave populations of *A. mexicanus*, though the number and sensitivity of responsive cells was severely reduced across cave populations. Imaging from the cavefish retina did not identify responses to flashes of light, raising the possibility that the light-induced activity that we observed in the tectum derives from internal photoreceptors and/or the pineal body (*64*). These differences support the hypothesis that the light response in the tectum is vestigial in cavefish, suggesting a transformation in the function of the cavefish tectum. Also in support of this hypothesis is the finding that fish from the evolutionarily older Pachón population exhibit more severe deficits in tectal light response relative to fish from the Molino caves.

In zebrafish, the tectum is primarily devoted to visual processing, but also receives mechanosensory and auditory information. Previous studies have identified a laminar organization of the zebrafish tectum, with different layers corresponding to distinct functional specializations (*27, 65, 66*). Numerous cellular response types have been previously identified in zebrafish, including those which respond selectively to changes in light level (*63*). Their identification in *A. mexicanus* provides an opportunity to investigate the genetic basis of tectal wiring through hybridization experiments. Our findings show that the prevalence of “On” cells and “Off” cells is uncorrelated in the tectum of hybrid individuals suggests that separate genetic mechanisms regulate these traits, likely distinct axon guidance molecules which regulate the wiring of the tectum (*67, 68*). Conversely, the correlation of both of these cell-types with dual-responsive “On/Off” cells, suggests a shared developmental basis, e.g. overlapping expression of axon guidance molecules. Further experiments in surface-cave F2 hybrids, in combination with quantitative trait locus (QTL) mapping, could elucidate the genetic basis of these, and other, functional cell types in the tectum.

Our analysis of the spatiotemporal correlations in the spontaneous activity within the tectum suggests cavefish have maintained functional connectivity despite the complete loss of a light-responsive retina and the near-complete loss of tectal light response. In the zebrafish optic tectum, spontaneous activity, reflects the functional connectivity of the circuit, which is adapted to improve visual detection of prey-like visual stimuli (*29*). These findings suggest that eye loss has not influenced the tectum’s functional connectivity. However, this trend did not hold true for the negative correlations within the tectum, which tended to be reduced in both Molino and Pachón populations. This suggests that tectal inhibition is tightly linked to visual processing, or that only positive correlations and not inhibition play a role in the potential repurposing of the optic tectum in cavefish.

It is possible that the visual circuitry of the sighted teleost has been repurposed for processing a different sensory modality in cavefish. This hypothesis is supported by the fact that in contrast to the significant reduction in visual response which may represent a vestige of visual degeneration evolution, the functional connectivity of surface and cavefish did not show significant differences. Systematic characterization of the response of the tectum to a battery of sensory stimuli can determine whether this brain region has been repurposed to respond predominantly to non-visual sensory stimuli.

The functional imaging described here represents the first use of Genetically Encoded Ca^2+^ Indicators to examine evolved differences in brain function in a vertebrate model. In addition to loss of vision, cavefish are widely used to study many different behaviors including enhanced lateral line function, sleep loss, prey capture, olfactory processing and stress (*11, 52, 69*). The application of whole-brain imaging approaches that have been developed in zebrafish to evolutionary models including *A. mexicanus* has the potential to identify neural mechanisms associated with evolved differences in these behaviors (*70*). Therefore, this manuscript provides a framework from applying comparative functional imaging to identify variability in brain function.

## Methods

### (a) Fish rearing and maintenance

Animal husbandry was carried out as previously described (*71*) and all protocols were approved by the IACUC Florida Atlantic University. Fish were housed in the Florida Atlantic University core facilities at 23 °C ± 1 °C constant water temperature throughout rearing for behavior experiments (*71*). Lights were kept on a 14:10 h light-dark cycle that remained constant throughout the animal’s lifetime. Adult fish were fed a diet of blood worms to satiation 2-3x daily (Aquatic Foods, Fresno, CA), and standard flake fish food during periods when fish were not being used for breeding (Tetramin Pro).

### (b) Functional Imaging

Prior to imaging experiments, larvae were paralyzed by 60 s of immersion in 0.5mg/ml mivacurium chloride, rinsed thoroughly with system water, and embedded in 2% low-melting point agarose (Thermo-Fisher, A9414), in clear plastic 35mm dishes (Nunc, Thermo-Fisher, 150680), with a strip of white LED lights (Lepro) affixed rostral to the larvae. The microscope was shielded to prevent unintended light stimulus. Following a minimum of 1 minute of recording to capture basal activity levels, a 30 second visual stimulus was applied. In follow-up experiments, multiple 30 second stimuli were applied, separated by additional 1 minute intervals. Visual stimulus was controlled by custom code running on Octave 6.2.0, utilizing the Arduino package. All visual stimulus experiments were acquired on a Nikon A1 confocal microscope, at 20X magnification, 2X digital zoom, at a rate of 1.96 Hz.

### (c) 2-photon calcium imaging

To capture spontaneous activity in the tectum, larvae were prepared as described above, and Ca^2+^ activity in the tectum was monitored for a period of 30 minutes, at a rate of 3.96 Hz on a Nikon A1R Multiphoton microscope, with a Chameleon Vision II Ti:Sapphire tunable laser at an excitation wavelength of 940 nm. Images were acquired with a 25x objective, and 1.5X zoom.

### (d) Ca^2+^ activity analysis

Ca^2+^ activity of individual cells was extracted using a MATLAB toolbox developed by Romano et al. (*38*). Briefly, an average image was generated from acquired time-lapse images, and a watershed algorithm was applied to generate ROIs outlining individual cells. These were used to extract raw fluorescence data from the time-lapse images, and calculate the ΔF/F values for individual cells.

### 1.1 (e) Correlation analysis

In order to compute the average probability density function (pdf) for correlations in surface, Molino and Pachón, we estimated the correlation pdf for each larva by counting the number of correlations in regularly spaced (0.05) correlation bins between −1 and 1. We then averaged those pdf together within each population. In order to quantify the amount of correlation that emerged from the spontaneous rate of activity, we built a null model by shifting circularly the neuron’s activity time series by a random lag between 0 and 30min. The latter corresponds to the total duration of the experiments. This keeps the time series untouched while destroying the temporal correlations between time series. We performed 1000 repetitions of this procedure to obtain the null model. To compare the distributions of correlations between the surface, Molino and Pachón populations, we quantified the distance between the null-model distribution and the distribution of correlations in our dataset using the Jensen-Shannon divergence for each larva. The discrete Kullback-Leibler divergence *D*_*KL*_, or relative entropy, quantifies the similarity between probability distributions *P* and *Q* defined on some probability space *X*.

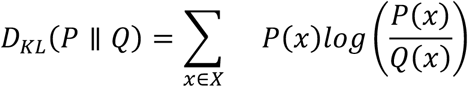

The Jensen-Shannon divergence *D*_*JS*_ is based on the Kullback-Leibler divergence, but is symmetric, such that *D*_*JS*_ (*P* ∥ *Q*) = *D*_*JS*_ (*Q* ∥ *P*). It can be defined as:

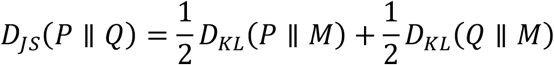

where

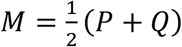

Finally, to investigate the spatial structure of correlations in the optic tectum of surface, Molino and Pachón, we averaged the correlation between pairs of neurons falling into regularly spaced (20 *µm*) distance bins between 0 and 200 *µm* to obtain the distribution of correlations against distance.

### 1.2 (f) Identification and categorization of cells responsive to whole-field visual stimuli using linear regression

To identify and categorize neurons that are responsive to visual stimuli, we used a multivariate linear regression model. To search for neurons with specific responding profiles, we used a regressor-based analysis. We used 3 binary regressors based on the timings of the onset and offset the visual stimuli (*R*_1_) or the offset (*R*_2_) of the visual stimulation, or a sustained activation or inhibition during the whole duration of the stimulus (*R*_3_). To model the slow dynamics of the H2B-GCaMP6s calcium sensor, we convolved those regressors with a decaying exponential kernel with characteristic time constant *τ* = 3.5s (*72*). More specifically, if we denote *y*_*n*_ the time series of the relative change of fluorescence over time for neuron *n*, we have:

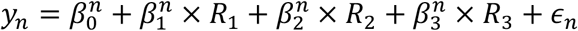

Where 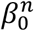 is the intercept, 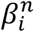 are the regression coefficients associated to each regressor *R*_*i*_, and *ϵ*_*n*_ represent the residuals for neuron *n*. We compared the fraction of variance explained by the linear model, *R*^2^, which is a measure of the goodness of fit and compared this to a null model. We also compared the regression coefficient *β*_*i*_ obtained for each regressor, which captured the intensity of the response of a neuron, against the distribution of regression coefficients under the null model.

A neuron was classified as responsive if its *R*^2^ value exceeded the 95^th^ percentile of the distribution of *R*^2^ over the whole population of recorded neurons in the null-model 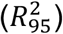, and if exactly one of its regression coefficient was also higher than the 95^th^ percentile of the null *β* distribution for this neuron 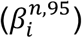, or lower than the 5^th^ percentile in the case of inhibition 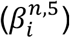. This procedure ensures that silent neurons, which may present a very good fit (high *R*^2^ values) but low *β* value are excluded from the population of responsive neurons, and also allows to categorize the neurons into 5 response profiles:

- ON cells: 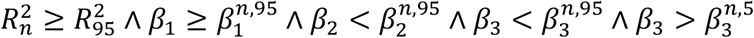
- OFF cells: 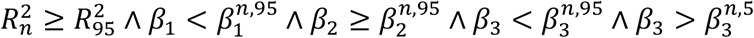
- ON/OFF cells: 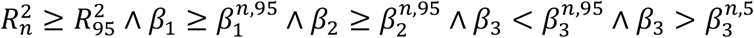
- Sustained cells: 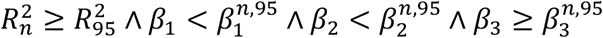
- Inhibited cells: 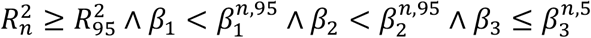

### 1.3 (g) Detection of significant calcium events

In order to detect significant calcium events in the fluorescence time series, we used the method described in Romano et al., 2015 (*29*). This enables us to compute the fraction of neurons active at any time point during the recording.

### 1.4 (h) Statistical inference and data analysis

Non parametric statistical tests were performed when the normality assumption could not be upheld. Data analysis was performed using custom scripts written in MATLAB (R2020a).

---

**Supplemental Figure 1.**
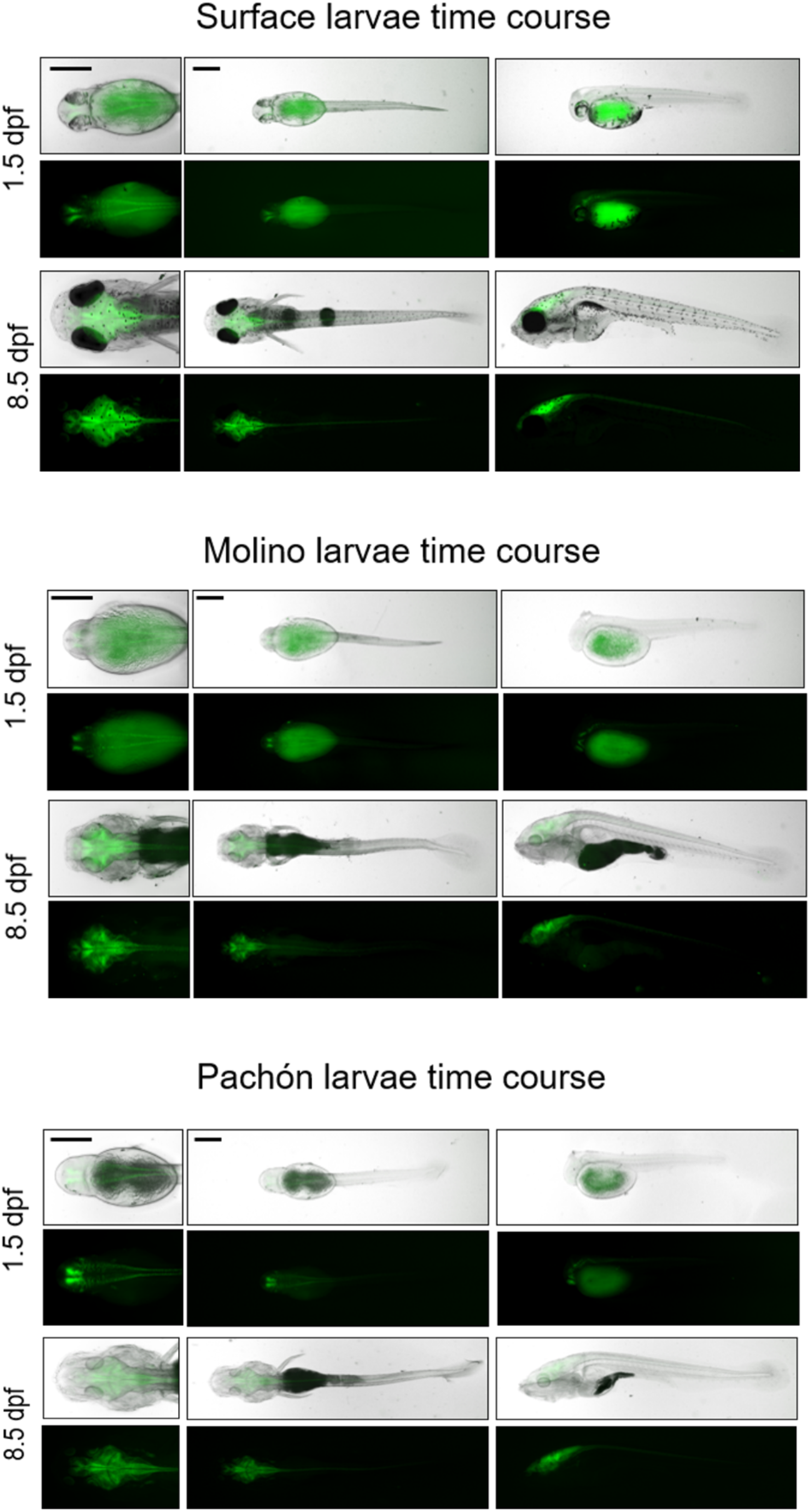
Stable expression of pan-neuronal HuC:GCaMP6s in surface and cave larvae. Brain-wide expression of HuC:GCaMP6s is detectable as early as 1.5dpf through at least 8.5dpf in surface **(A)**, Molino **(B)**, and Pachón **(C)** larvae.

**Supplemental Figure 2.**
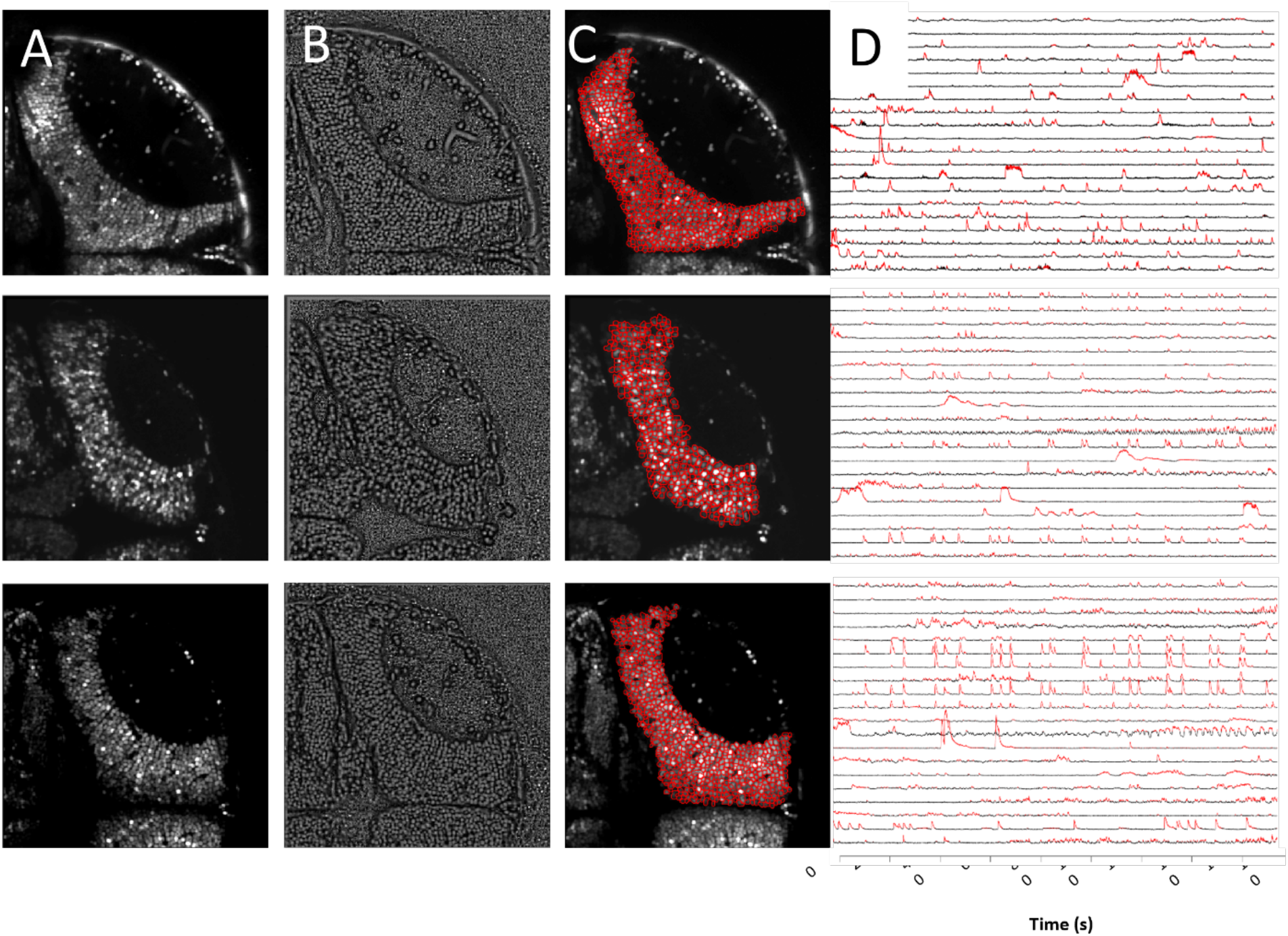
Automated segmentation of neuronal ROIs. **(A)** Representative images of the superficial tectum in surface fish (top), and fish from the Molino (middle) and Pachón (bottom) caves, harboring pan-neuronal HuC:GCaMP6s. **(B)** Result of local contrast normalization to enhance boundaries between cells, preparing the image for segmentation. **(C)** Result of watershed segmentation, with individual cells outlined in red. **(D)** Representative subset of Ca^2+^ traces of individual neurons obtained through automated segmentation. Scale bar = 50 microns

**Supplemental Figure 3.**
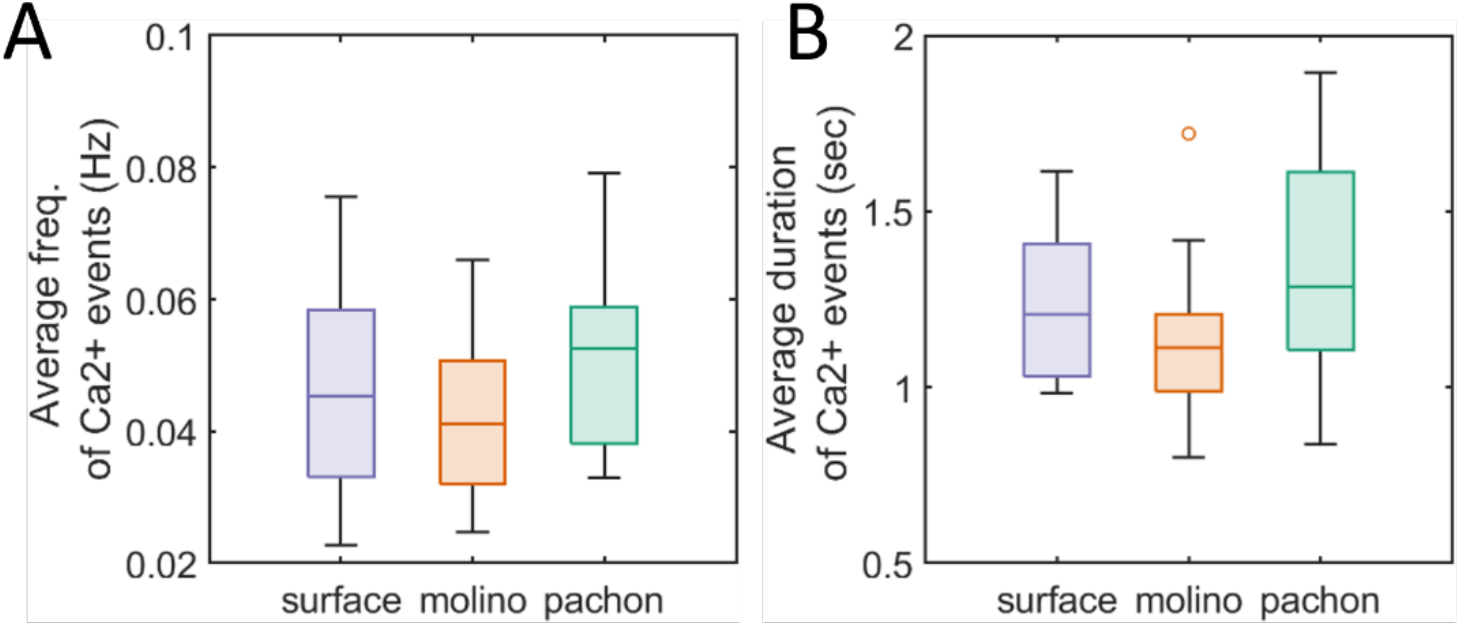
Maintenance of tectal activity in blind cavefish. There is no significant difference in the frequency (Surface: 0.047 ± 0.018 Hz, Molino: 0.042 ± 0.011 Hz, and Pachón: 0.051 ± 0.014 Hz, mean ± standard deviation, one-way ANOVA) **(A)** or duration (Surface: 1.246 ± 0.226 s, Molino: 1.123 ± 0.210 s, and Pachón: 1.324 ± 0.325 s, mean ± standard deviation, one-way ANOVA). **(B)** of Ca^2+^ events in the tectum of blind cavefish from the Molino (orange) or Pachón (green) caves.

**Supplemental Figure 4.**
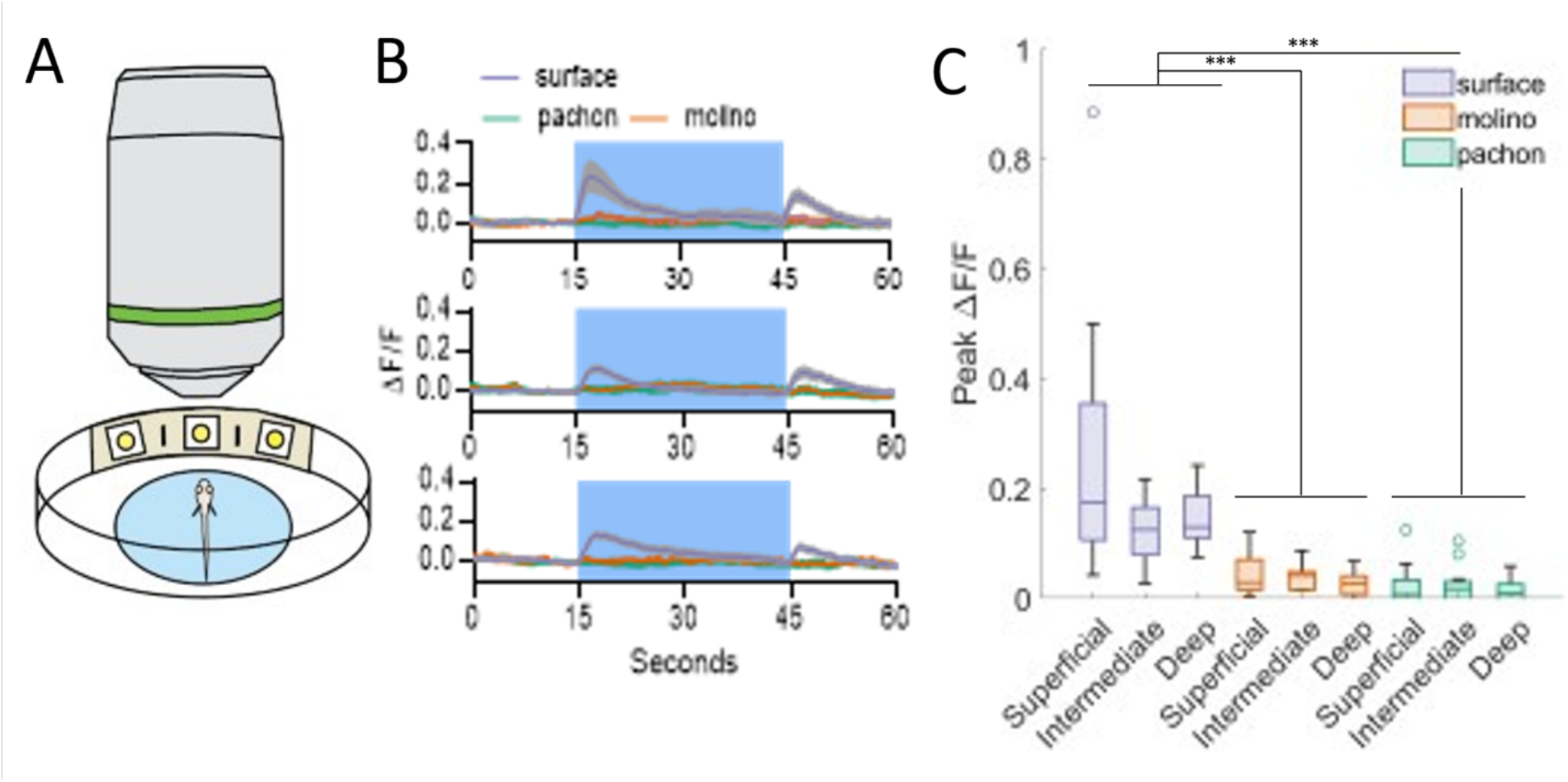
Loss of response to visual stimulus in the cavefish optic tectum. **(A)** Scheme of the system used for visual stimulation showing the larva embedded in agarose, the 3 LEDs and the objective. **(B)** Mean ΔF/F values in the tectum before, during, and after 30 seconds of white light presentation, in the superficial (top), intermediate (middle), and deep (bottom) tectum. **(C)** Peak ΔF/F values during 30 seconds of white light stimulus, at three depths within the tectum (two-way ANOVA, *F*_4,112_ = 3.093, *p*=0.02).

**Supplemental Figure 5:**
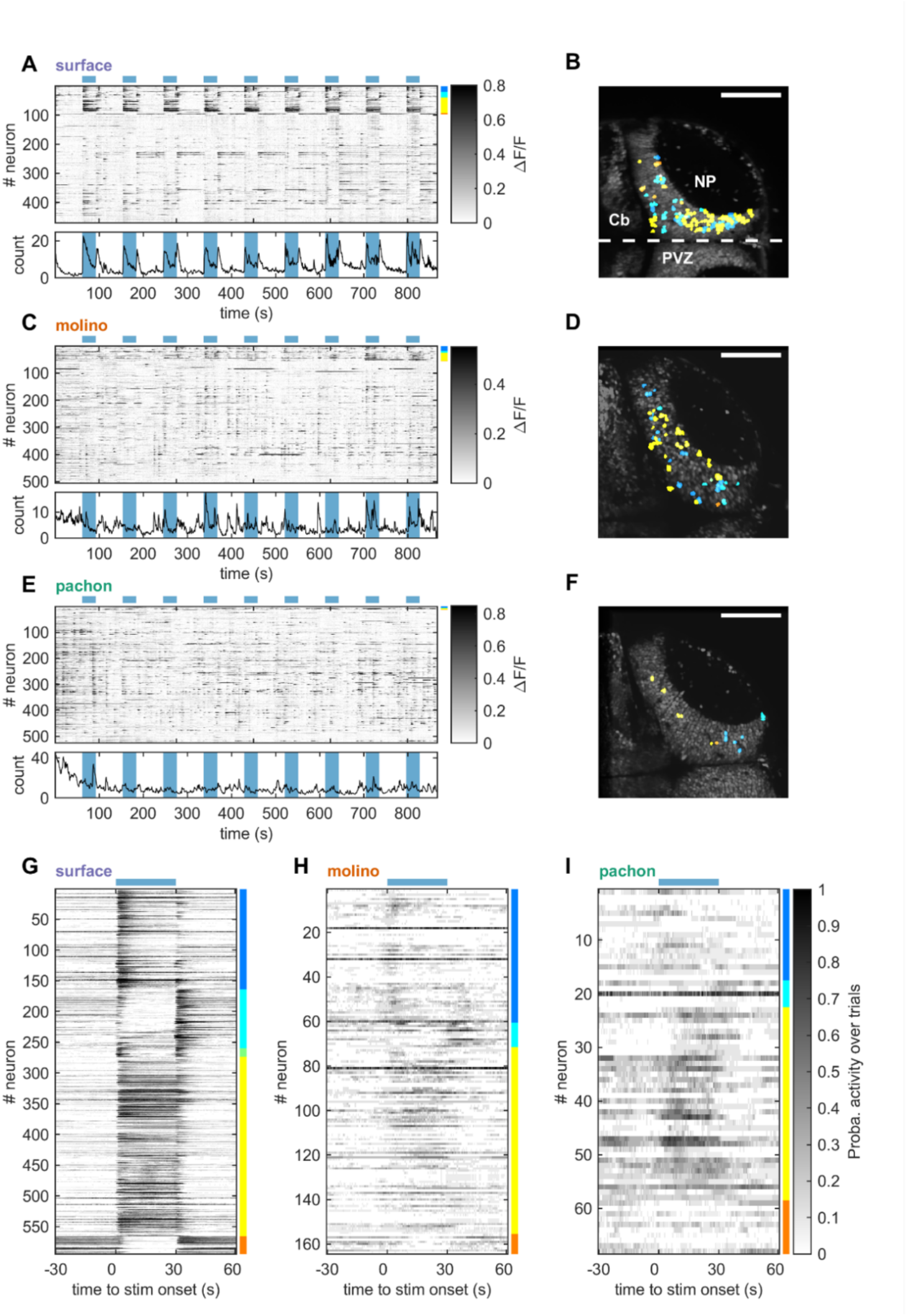
Neuronal responses to whole-field visual stimuli. **(A**,**B**,**C)** Top: Representative raster plot of neuronal activity in response to whole-field visual stimuli for single surface (A), Molino (B) and Pachón (C) larvae. Visual stimuli were delivered by an array of white LEDs for a duration of 30 seconds (represented by horizontal light blue bar above the raster), with an inter-stimulus interval of 120 seconds, and neuronal activity was recorded with a frame rate of 1.96 Hz. Each experiment contains between 8 and 20 visual stimuli. Neurons responsive to the visual stimuli were detected using a linear regression model and classified into 5 categories: “on” cells (blue), “off” cells (turquoise), “on/off” cells (green), “sustained” cells (yellow) and “inhibited” cells (orange). The raster was sorted with responsive neurons at the top. Bottom: proportion of significant calcium events. Shaded light blue regions denote the presentation of visual stimuli. **(D**,**E**,**F)** Topography of responsive neurons corresponding to the raster plots in (A,B,C). NP: neuropile, Cb: cerebellum, PVZ: peri-ventricular zone. The dotted white line represents the midline of the brain. Scale bar: 100 μm. **(G**,**H**,**I)** Raster plots of the probability of occurrence of significant calcium transient around whole-field visual stimulation (horizontal light blue bar) across stimuli repetitions, over the whole population of surface (A, n=6), Molino (B, n=5), and Pachón (C, n=3) larvae.

**Supplemental Figure 6.**
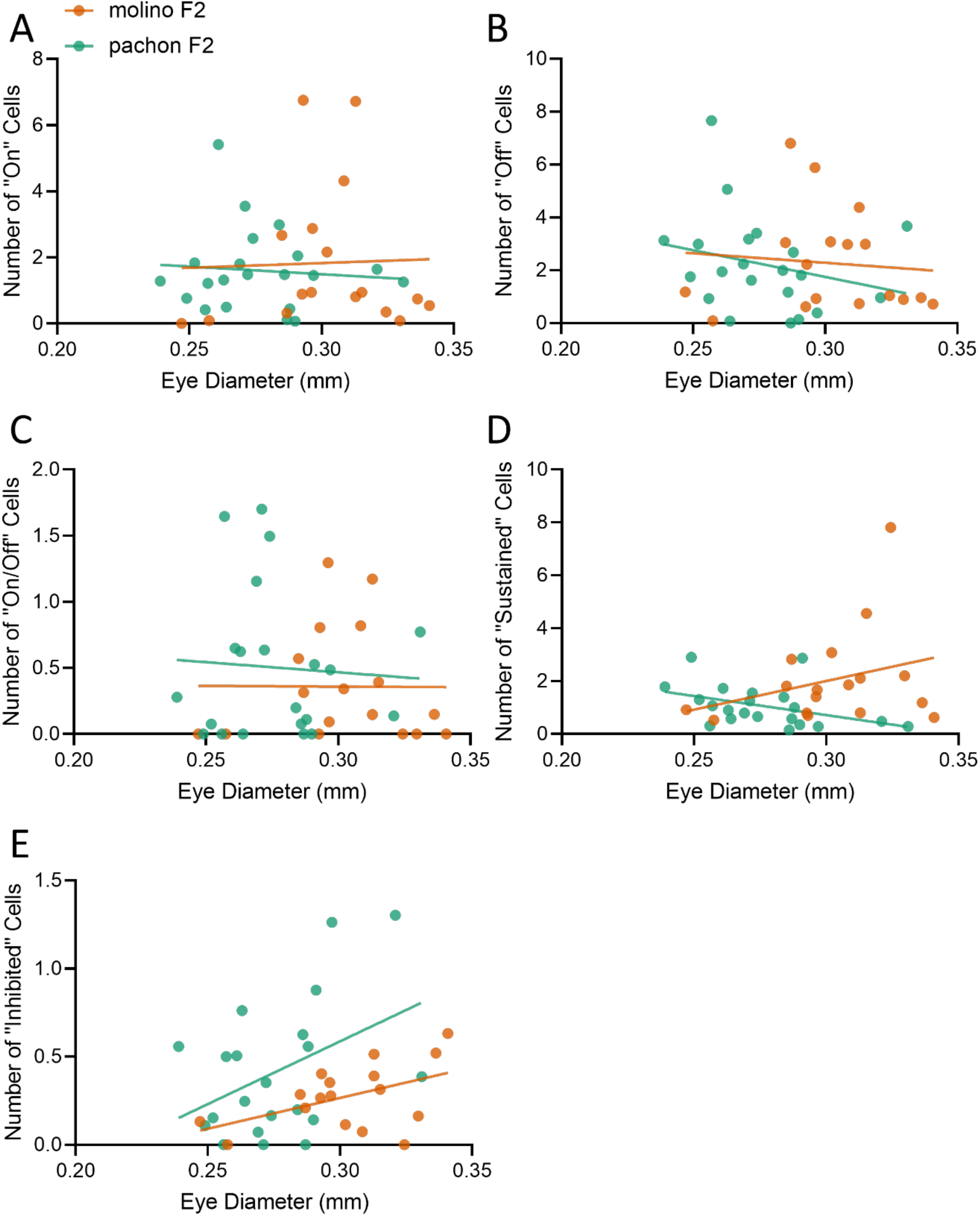
Eye size is not correlated with light response in surface-cave hybrids. There is no significant correlation between eye diameter and the proportion of cells responding to light stimulus, in any of the cell types identified, in either Molino (orange) or Pachón (green) F2 hybrids. (A) “On”: Molino F2 (*r*^2^=0.001, *F*_1,15_ = 0.016, *p*=0.9), Pachón F2 (*r*^2^<0.001, *F*_1,19_ = 0.137, *p*=0.71). (B) “Off”: Molino F2 (*r*^2^=0.008, *F*_1,15_ = 0.131, *p*=0.72), Pachón F2 (*r*^2^=0.07, *F*_1,19_ = 1.312, *p*=0.26). (C) “On/Off”: Molino F2 (*r*^2^=0.003, *F*_1,15_= 0.074, *p*=0.79), Pachón F2 (*r*^2^<0.001, *F*_1,19_ = 0.001, *p*=0.98). (D) “Sustained”: Molino F2 (*r*^2^=0.09, *F*_1,15_ = 1.478, *p*=0.25), Pachón F2 (*r*^2^=0.18, *F*_1,19_ = 4.083, *p*=0.06). (E) “Inhibited”: Molino F2 (*r*^2^=0.23, *F*_1,15_ = 4.506, *p*=0.051), Pachón F2 (*r*^2^=0.18, *F*_1,19_= 4.128, *p*=0.056).

